# Tissue-specific and functional loci analysis of *CASP14* gene in the sheep horn

**DOI:** 10.1101/2024.09.03.611085

**Authors:** Xiaoning Lu, Guoqing Zhang, Hao Yang, Mingzhu Shan, Xiaoxu Zhang, Yuqin Wang, Junyan Bai, Zhangyuan Pan

## Abstract

Under the current context of intensive farming, small-horned animals are more suitable for large-scale breeding. The *CASP14* gene, closely associated with skin and keratin formation, may influence horn size due to its link with skin development. This study comprehensively analyzed the tissue-specific expression of *CASP14* using RNA-Seq data, identified functional sites through whole-genome sequencing (WGS), and investigated allele-specific expression (ASE) validated by KASP assays. Results showed significantly higher *CASP14* expression in the scurred group com-pared to the SHE group, with pronounced expression in the skin. Interbreed comparisons also revealed elevated *CASP14* levels in the scurred group. Analysis of potential functional sites indicated structural similarities in the *CASP14* protein among horned mammals. Five ASE events were discovered, and intersecting these with SNPs and high fixation index loci (Fst > 0.05) identified three potential functional sites: 7941628, 7941817, and 7941830. The SNP site 7944295 was selected for T-test analysis and further validated through KASP assays, corroborating the role of *CASP14* in sheep horn phenotypes. Our findings suggest that *CASP14* plays a significant role in horn development, offering insights into breeding strategies for small-horned animals.

## Introduction

Sheep is a significant livestock species, playing a crucial role in the development of the national economy and the supply of meat[1]. To enhance the production efficiency and productivity of sheep through genetic improvement techniques, understanding traits such as growth, body weight, carcass quality, wool quality, and horn type is essential [2, 3]. Animal horns are considered cranial appendages, predominantly found among ruminant species [4]. Horns are distinctive cranial appendages of the *Bovidae* family, which includes species such as *Bos taurus (cattle), Ovis aries (sheep)*, and *Capra hircus (goat)* [5]. To safeguard animals and producers from accidental injuries, polled animals better meet the requirements of current intensive livestock management practices [6-8]. Therefore, sheep horns serve as an important model for the study of phenotypic genetic evolution in animals [9]. However, the majority of current research on sheep focuses on the genetic mechanisms of their primary eco-nomic traits [10]. The genetic mechanisms underlying sheep horn development remain unclear.

Caspases (*CASP*) are cysteine proteases that play a central role in apoptosis. Therefore, they may also be involved in the terminal differentiation of keratinocytes [11]. Caspase-14 (*CASP14*) is a member of the *CASP* family. *CASP14* is a cysteine-dependent aspartate-specific protease that is expressed during the process of epidermal differentiation, is highly expressed in the skin of mammals. In humans, this protein is specifically localized to the differentiating keratinocytes, suggesting its critical involvement in the development of the epidermal barrier [12]. *CASP14* is involved in the process of keratinocyte apoptosis [11]. Furthermore, the expression of *CASP14* mRNA is confined to the uppermost viable layers of the epidermis, encompassing the granular layer, hair follicles, and sebaceous glands. This distribution implies that *CASP14* may have broader functions beyond its role as a proapoptotic gene [13]. Studies on BeWo cell lines, a choriocarcinoma-derived model for trophoblasts, have revealed a contrasting role of *CASP14* compared to its function in human epi-dermis. Inhibition of *CASP14* in these cells resulted in the upregulation of KLF4, hCG, and cytokeratin-18—markers associated with normal trophoblast differentiation. These findings suggest that *CASP14* influences the differentiation pathway of trophoblasts [14, 15]. Another research has indicated that the *CASP14* gene exhibits variations associated with skin immune responses and diseases, as well as variations related to maintaining skin immune homeostasis following chemical exposure [16]. Thus, the *CASP14* gene plays a significant role in the formation of the skin barrier in animals.

Base on the previous research of our team, we have identified differentially expressed genes between large and small sheep horns and conducted Gene Ontology (GO) enrichment analysis. The results suggest that the *CASP14* gene is involved in processes such as skin barrier formation. Horns are derivatives of the skin and represent an independent organ, the development of the stratum corneum is believed to be primarily regulated by the skin [17], we speculate that the *CASP14* gene also plays a significant role in the formation of sheep horns. To further validate this hypothesis, we initially utilized RNA-sequencing (RNA-Seq) data to explore the tissue-specific expression of the *CASP14* gene to understand its function. Subsequently, we employed whole-genome sequencing (WGS) data to analyze functional loci within the *CASP14* gene associated with sheep horns. This study aims to investigate the molecular mechanisms underlying the formation of sheep horns, providing valuable molecular markers for sheep horn breeding.

## Materials and Methods

### Ethics

Institutional Review Board Statement: All the experimental procedures mentioned in the present study were approved by the Science Research Department (in charge of animal welfare issues) of the Institute of Animal Sciences, Chinese Academy of Agricultural Sciences (IAS-CAAS) (Beijing, China). Ethical approval on animal survival was given by the animal ethics committee of IAS-CAAS (No. IASCAAS-AE-03, 12 December 2016).

### Sample Collection

All RNA-Seq data were obtained from our laboratory (PRJNA1003277) [18], totaling fifteen Tibetan sheep samples from Dangxiong, Tibet, China. All individuals in this study are female, with ages ranging from 2 to 4.5 years. The Tibetan sheep were categorized into two groups based on horn characteristics: one with 7 sheep having scurred horns (0–12 cm), and another with 8 sheep with SHE horns (>12 cm). From the 15 sheep, we collected tissue with soft horns, placed them in deep cryopreservation tubes, and stored the tubes in liquid nitrogen.

### RNA Sequencing Data Processing

Our data was sourced from publicly available and laboratory collections. A total of 2,915 high-quality RNA-Seq datasets were accessed from the National Center for Biotechnology Information (NCBI) and the European Bioinformatics Institute (EBI). The data were processed alongside laboratory-collected samples through the following procedures. Processing and removal of low-quality bases and artifact sequences were implemented using the TrimGalore (v.0.6.7). The clean reads were aligned to the sheep reference genome ARS-UI_Ramb_v2.0, using the STAR (v.2.5.4) with the “--chimSegmentMin 10” and “--outFilterMismatchNmax 3” parameters. High-quality RNA-Seq clean datasets were acquired for subsequent analysis, characterized by unique mapping reads exceeding 85% and a count of clean reads greater than 20,000,000. Furthermore, gene expression levels were standardized using StringTie (v.2.1.5) by calculating fragments per kilobase of transcript per million mapped reads (FPKM) and transcripts per million (TPM). Ultimately, the featureCounts (v.2.0.1) was utilized to extract the raw counts of genes.

### Analysis of *CASP14* Gene Expression

To explore the difference in *CASP14* gene expression between scurred and SHE groups, a boxplot was generated using the ggplot2 package (v.3.4.4) in R (v.4.3.0). To further analyze the distinctions in *CASP14* exon expression between these two groups, the genome annotation (GTF) file was formatted using the dex-seq_prepare_annotation2.py script from the Subread_to_DEXSeq package (v.1.46.0). The formatted GTF file and the counts matrix produced by featureCounts were processed using load_SubreadOutput.R in RStudio, enabling the construction of the DEXSeqDataSetFromFeatureCounts (dds) object. Subsequently, an exon difference analysis was conducted to identify differential exon expression between the groups.

We obtained RNA-Seq data from the Genotype-Tissue Expression (GTEx) project, encompassing a total of 2651 pig samples, 4359 cow samples, and 9810 human samples, along with their TPM values. RNA-Seq data from sheep, pigs, cows, and humans were aggregated and utilized to categorize samples into 16 tissue types, subsequently computed the average TPM values for each species. Moreover, data on eight sheep breeds were extracted from public RNA-Seq databases, the selected breeds were Carpet, Rambouillet, Tibetan, Bashibai, Chinese Merino, Hu, Spanish Churra, Tan, which were categorized into scurred, and SHE subgroups.

### Evolutionary and Structural Analysis of *CASP14*

We retrieved the FASTA files for the gene of interest in 20 different species from the NCBI database and constructed a phylogenetic tree for the *CASP14* gene using the online tool iTOL (https://itol.embl.de/). To ascertain whether the amino acids are located at key positions, we predicted the 3D structure of the *CASP14* protein using AlphaFold2 (https://colab.research.google.com/github/sokrypton/ColabFold/blob/main/AlphaFold2.ipynb), which generated five models. From these, we selected the model with the highest score.

### Analysis and Validation of Whole-Genome Sequencing

The sequencing data sets consisted of WGS generated in this study and published previously, including 3125 sheep from PRJNA304478, PRJNA325682, PRJNA479525, PRJNA624020, PRJNA675420, PRJNA822017, PRJNA30931, PRJNA480684, PRJNA509694, PRJNA779188, and PRJNA783661 [19-30]. A sample of 3,125 sheep was classified into scurred and SHE groups. F-statistic (Fst) values were calculated using VCFtools (v.0.1.16) to identify SNP loci exhibiting significant differences between these two populations. Small Tail Han sheep were selected as the experimental breed, and horn length was quantified in 36 individuals by measuring from the horn base to its outermost end. The right horn length was used to determine the overall horn size. Sequencing reads were trimmed using Trimmomatic (v.0.39), and the raw sequencing data quality was assessed with FastQC (v.0.12.1). The qualified reads were then aligned, sorted, and merged to the sheep reference genome using BWA (v.0.7.17) and Picard (v.3.1.1). Mutation information was annotated using SnpEff (v.4.3). A dominant model was applied to establish the relationship between horn length and different genotypes. For further analysis of the SNP chain, the VCFtools (v.0.1.16) filters --maf 0.45 and --min-meanDP 5 were used, and LDBlockShow (v.1.40) was employed to identify SNP loci. Finally, the identified loci were sent to a biotechnology company for validation through Kompetitive Allele-Specific PCR (KASP) assays.

## Results and discussion

### Comparative *CASP14* Expression Across Sheep Horn Phenotypes

The expression level of the *CASP14* gene was significantly higher in the scurred compared to the SHE (p=0.019), as shown in Fig 1A. Subsequently, in Fig 1B, the expression of *CASP14* gene exons was found to be higher in the scurred. As shown in Fig 1C, all exons of the *CASP14* gene are expressed in both the scurred and SHE groups. The *CASP14* gene exhibits relatively high expression in the skin, while its ex-pression levels are low or even undetectable in the blood, CNS, heart, liver, lung and muscle.

**Fig 1.**
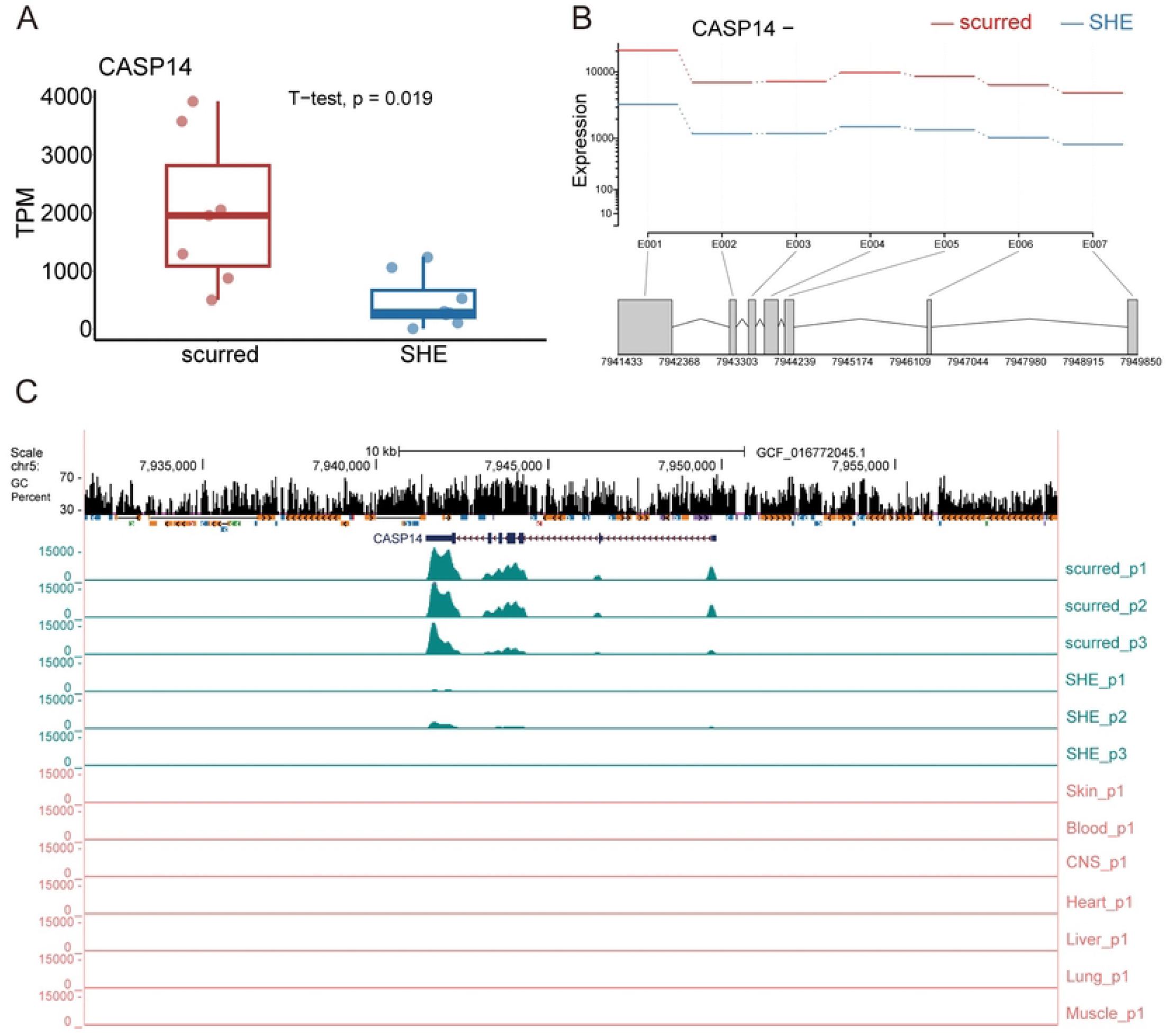
Expression of *CASP14* Gene in Scurred and SHE Groups. (A) Differential expression of *CASP14* gene between scurred and SHE Groups. (B) Expression of *CASP14* exons in scurred and SHE groups. The expression refers to the estimated values derived from the fitted expressions in the GLM regression, where E001 to E007 represent the number of exons. (C) Differential expression of *CASP14* in horn and other tissues.

### *CASP14* Tissue Specificity in Sheep

Fig 2A demonstrates that the *CASP14* gene exhibits the highest expression in human skin, with sheep and cattle displaying the next highest levels. Further analysis assessed potential sex-related variations in *CASP14* gene expression across various sheep tissues. The results are illustrated in Fig 2B, which presents the comparative expression levels of the *CASP14* gene across different tissues in male and female sheep. *CASP14* gene expression in skin tissue is elevated in both female and male subjects relative to other tissues. Additionally, the expression of the *CASP14* gene was examined across different species. In Fig 2C, among the three breeds of the SHE group (Carpet, Rambouillet, Tibetan), *CASP14* gene expression was the lowest. Conversely, higher expression levels were noted in scurred sheep breeds (Bashibai, Chinese Merino, Hu, Spanish Churra, Tan). These findings suggest that the *CASP14* gene is highly expressed in skin tissues across multiple species, with no significant differences in expression between females and males. Moreover, variations in expression levels were observed between the scurred and SHE group.

**Fig 2.**
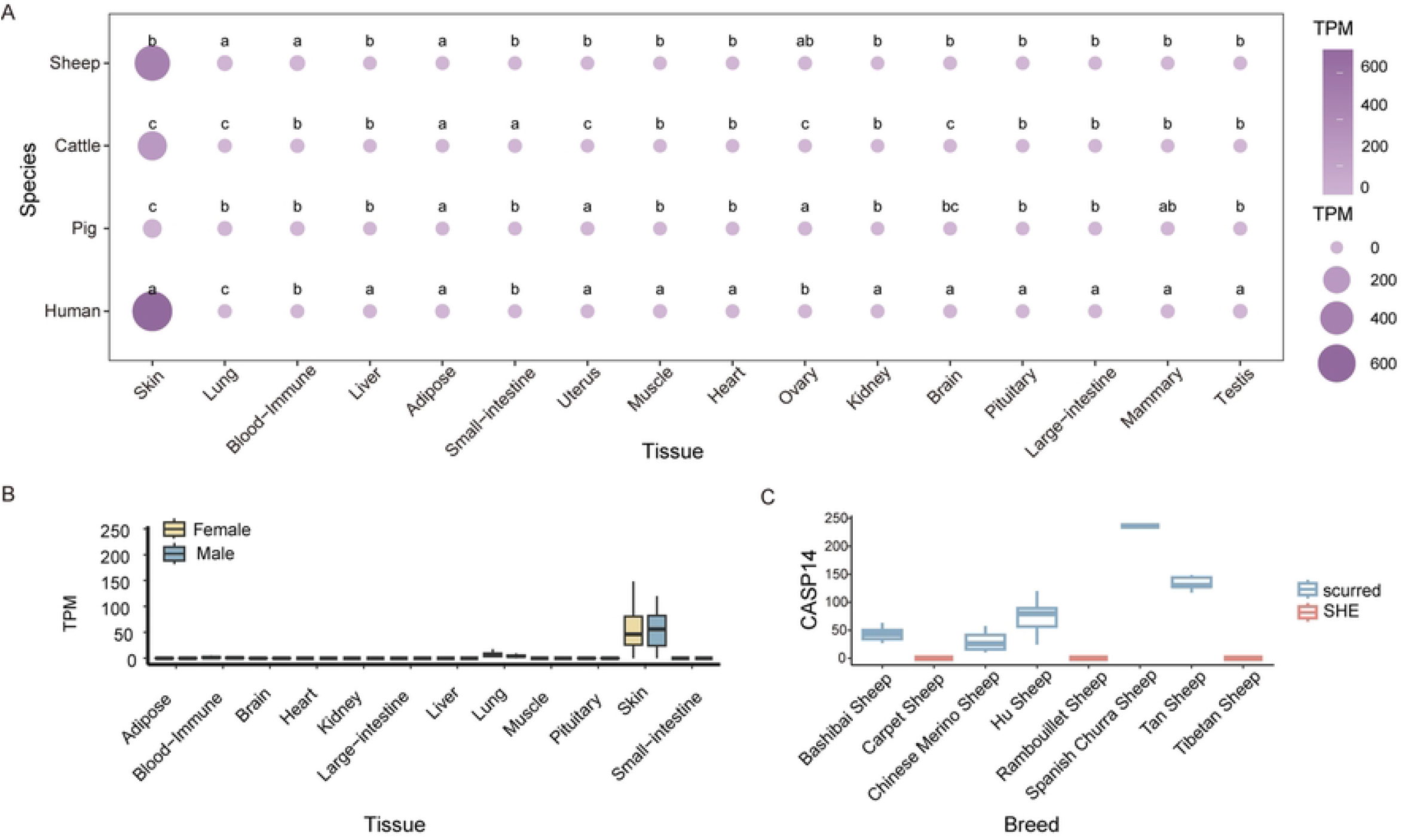
Expression Differences of *CASP14* Gene. (A) Differential expression of *CASP14* gene in four species. (B) Sex differences in *CASP14* gene expression across various sheep tissues. (C) Ex-pression of *CASP14* gene in various sheep breeds.

### The potential functions of the *CASP14* gene

As depicted in Fig 3A, the phylogenetic analysis of the *CASP14* gene aligns with the evolutionary relationships among species, showing that *Bovidae* and *Cervidae* cluster together. This indicates structural similarities in the *CASP14* protein among these horned mammals. Fig 3C illustrates amino acid sites unique to horned animals, with positions 11, 146, 148, 152, 153, 169, 172, 198, and 218 exhibiting a consistent pattern. These conserved residues may play a crucial role in the growth and development of sheep horns, indicating their potential functional importance in horn morphogenesis. Furthermore, as shown in Fig 3B, the protein structure of *CASP14* reveals that the amino acid at position 218, situated within a beta-sheet region, is consistently methionine in horned animals such as sheep, suggesting a significant association with horn presence.

**Fig 3.**
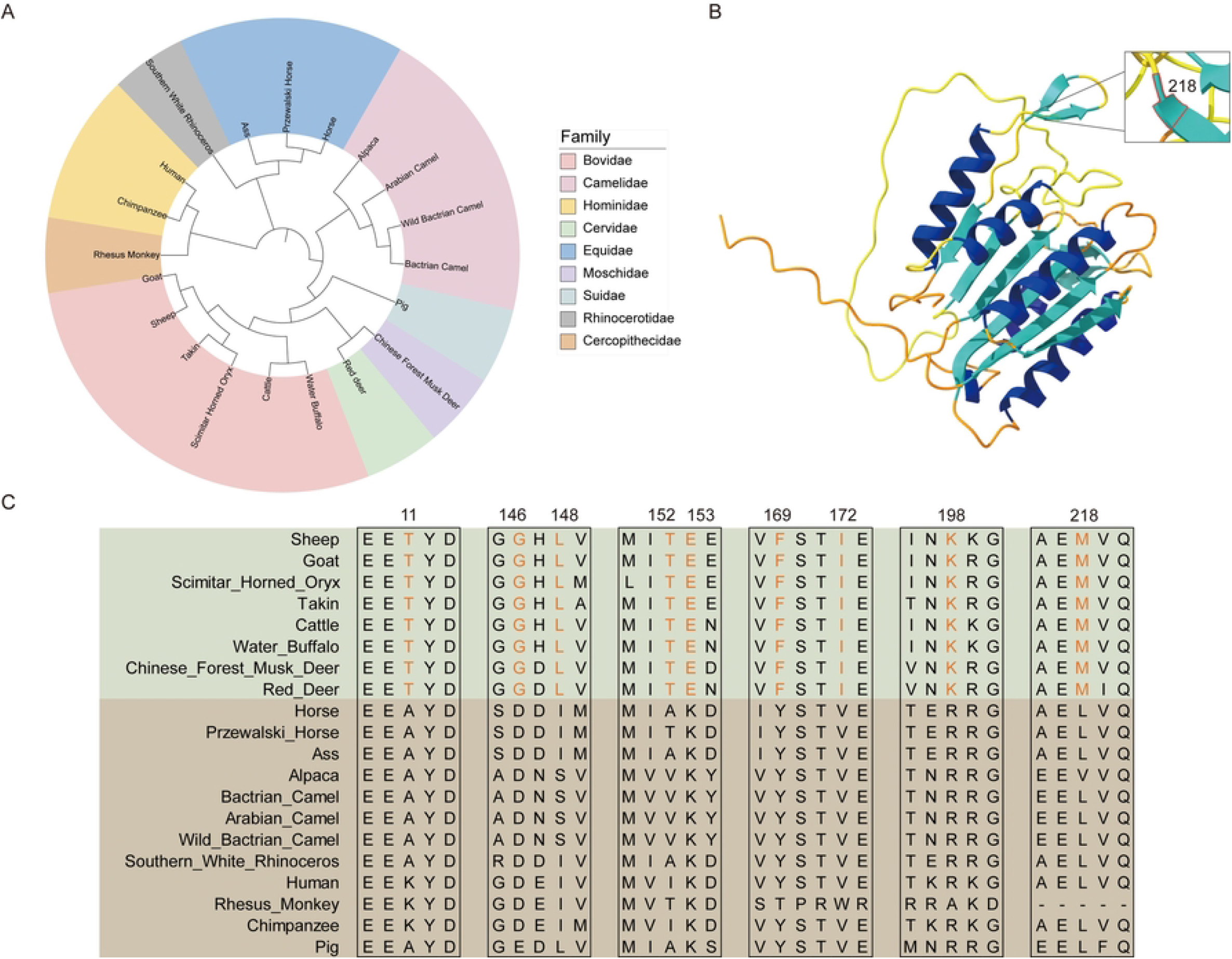
Structural and Evolutionary Insights of the *CASP14* Gene. (A) Evolutionary tree of the *CASP14* gene. (B) Three-Dimensional (3D) structure of sheep *CASP14* protein. (C) Amino acid sites of *CASP14* gene unique to horned animals.

### Characterization of Allele-Specific Expression in *CASP14* gene

As shown in Fig 4, analysis of RNA-Seq data revealed five allele-specific expressions (ASEs): Ase1 (Chr5: 7941514), Ase2 (Chr5: 7941569), Ase3 (Chr5: 7941628), Ase4 (Chr5: 7941817), and Ase5 (Chr5: 7941830). These ASE loci, all situated within Exon 7, might be closely linked to sheep horn types.

**Fig 4.**
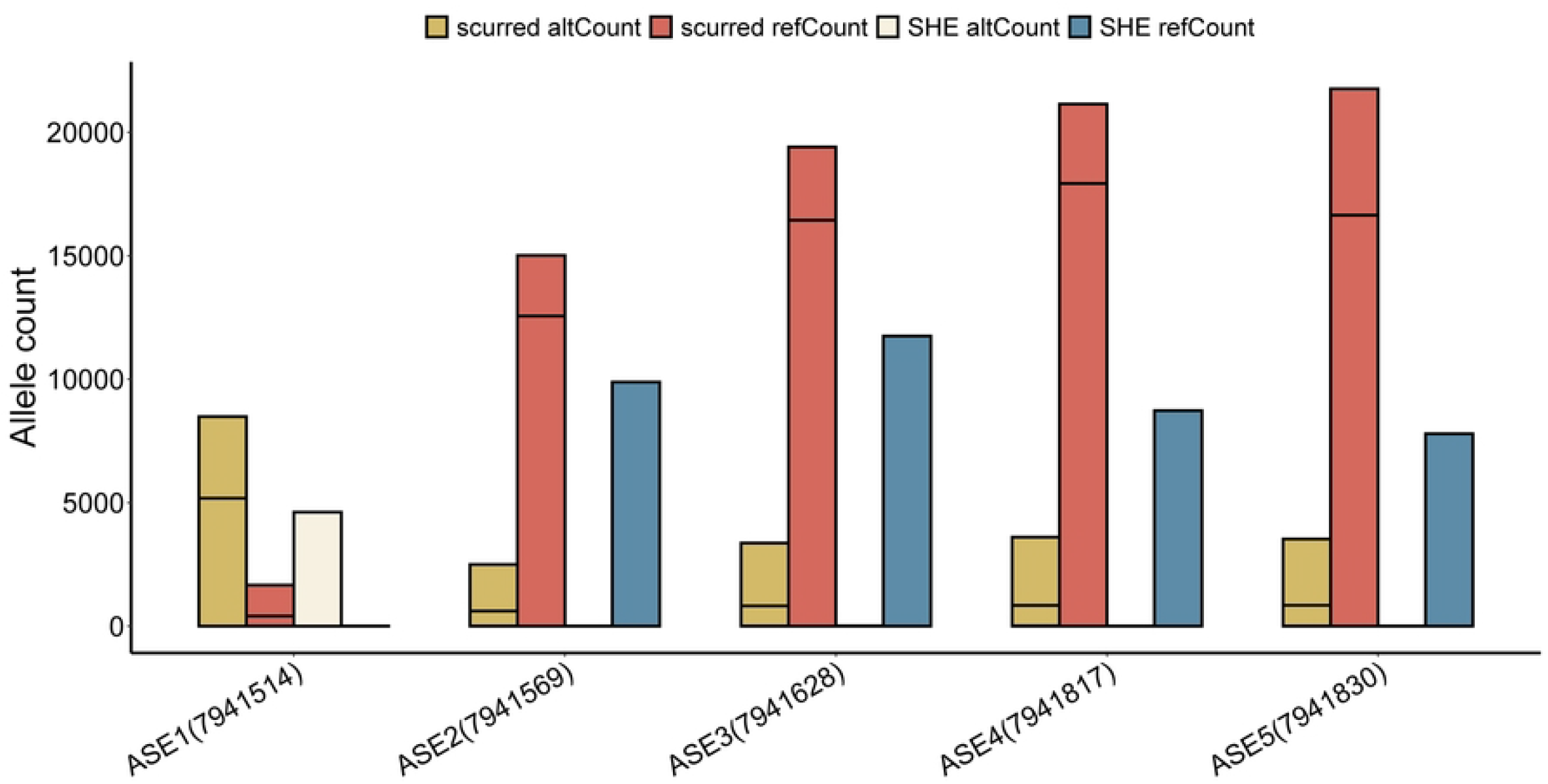
Allele count of 5 ASEs differing between scurred and SHE groups

### Potential Functional Mutations in the *CASP14* Gene

To observe the phenotypic differences between the scurred group and the SHE group under the influence of the *CASP14* gene, we conducted a principal component analysis (PCA). The results indicate a clear separation between scurred and SHE sheep breeds (Fig 5A). This suggests that the *CASP14* gene can differentiate between these populations, with loci within this gene region playing a regulatory role in horn development. Fig 5B identifies several loci with Fst values greater than 0.05, some of which are located within the exon regions of the *CASP14* gene. These loci have significant implications for studying the functional and adaptive differences of the *CASP14* gene between the scurred and the SHE populations. As depicted in Table 1, these are some SNP sites of the *CASP14* gene. Combined with the analysis of Fst values, these sites may be closely related to the size of sheep horns.

**Table 1.** Potential SNP functional loci of CASP14 gene.

| INFO | Mutation Type | Position | Fst | Ref | Alt |
| --- | --- | --- | --- | --- | --- |
| SNP1 | downstream | 7940550 | 0.0340909 | A | C |
| SNP2 | downstream | 7941388 | 0.0914412 | G | A |
| SNP3 | missense | 7943635 | 0.0602477 | T | C |
| SNP4 | intron | 7944295 | 0.0569244 | G | A |
| SNP5 | intron | 7946417 | 0.101679 | C | T |
| SNP6 | intron | 7946876 | 0.0784427 | G | A |
| SNP7 | exon | 7947041 | 0.0448705 | G | A |
| SNP8 | exon | 7947467 | 0.0181818 | C | T |
| SNP9 | upstream | 7950567 | 0.0151515 | C | T |
| SNP10 | upstream | 7950494 | 0.010101 | G | A |

**Fig 5.**
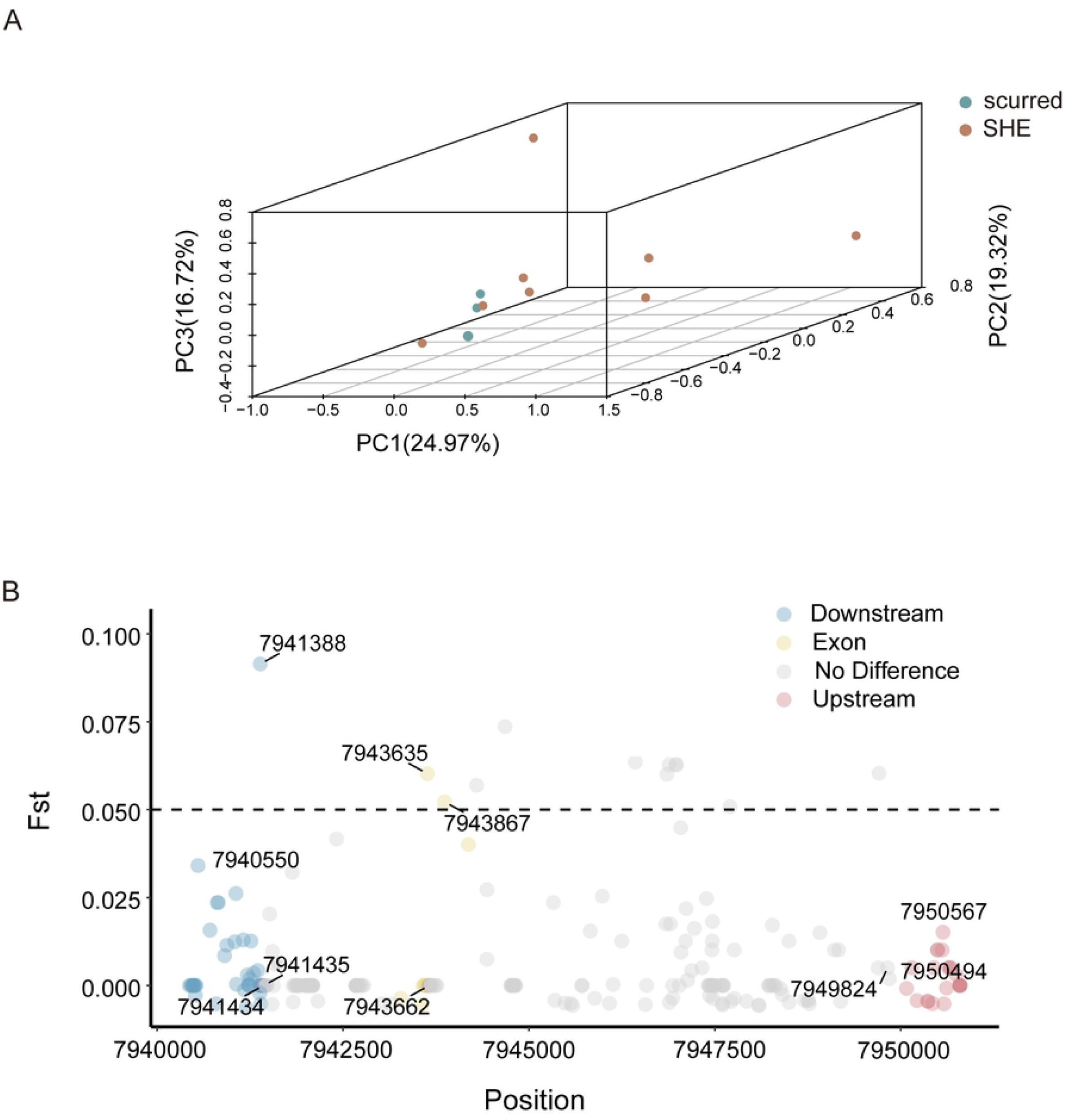
Genetic Variation and Population Structure of the *CASP14* Gene. (A) Three-dimensional (3D) principal component analysis (PCA) of the *CASP14* gene. Each point represents a sheep breed, which is categorized into scurred and SHE types based on breed information. (B) F-statistics (Fst) of functional loci of the *CASP14* gene.

In Fig 6A, the intersection of single nucleotide polymorphisms (SNPs), allele-specific expressions (ASEs), and Fst at the loci 7941628, 7941817, and 7941830, underscoring their critical roles in genetic diversity, population stratification, and gene expression modulation. These loci may represent genes or regulatory domains of functional significance, subject to selective forces, and implicated in the regulation of ASEs. Fig 6B illustrates the complex genetic associations among SNP loci within the 7.942 Mb to 7.950 Mb region on chromosome 5. Multiple high linkage disequilibrium (LD) blocks have been identified, indicating potential regions of functional importance where SNPs exhibit significant genetic linkage. The deep orange areas and SNPs within the black triangular regions, representing high LD, are focal points for further research and may contain genes related to important traits or diseases.

**Fig 6.**
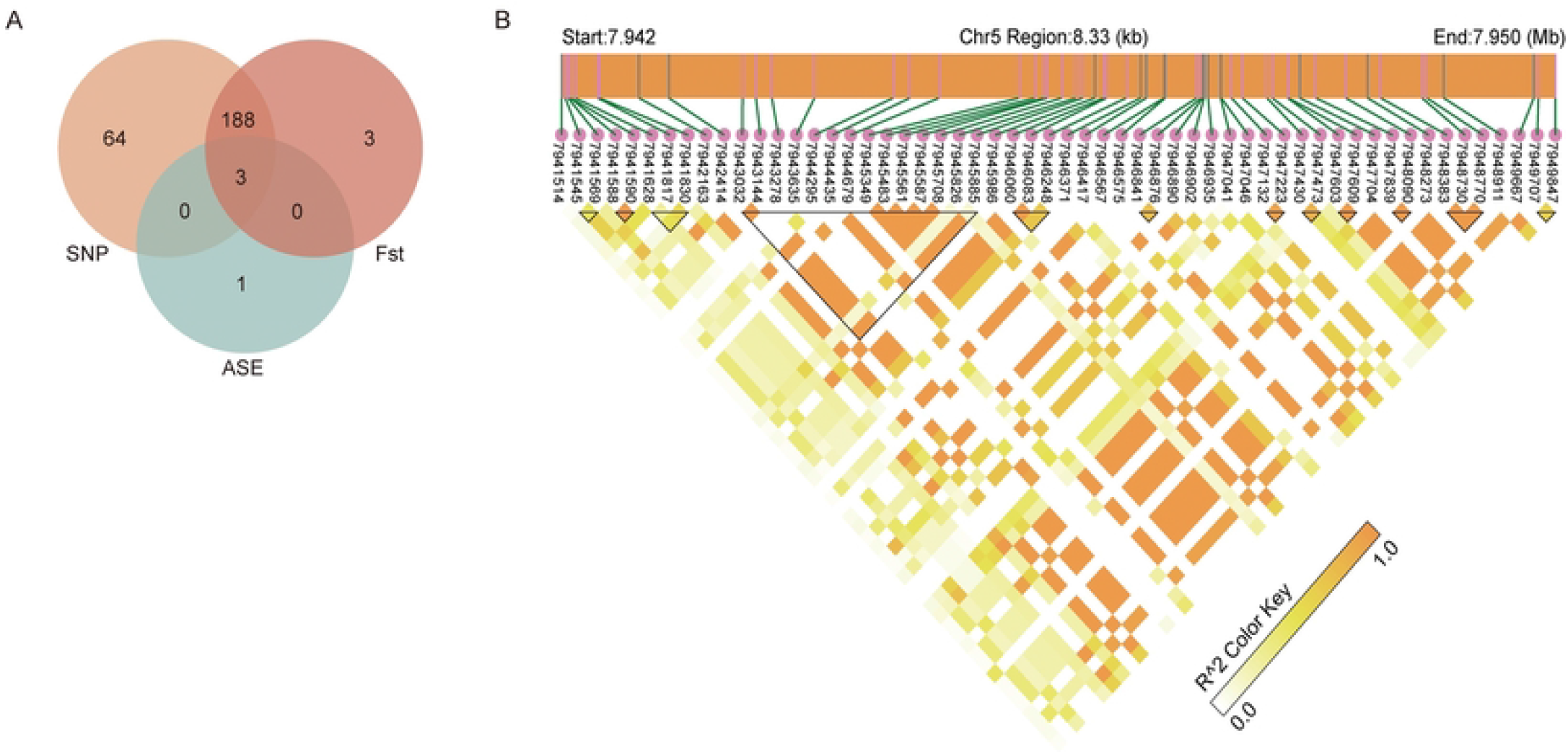
Genomic Insights into the *CASP14* Gene. (A) Intersections of ASEs, SNPs and Fst. (B) LD heatmap of *CASP14* gene. Darker colors represent higher LD values, and black triangles denote LD blocks, which are collections of SNPs with higher LD values.

### Validation of SNP Loci Using KASP Assay

Fig 7A illustrates a significant disparity in horn length among sheep with different genotypes at the specific loci chr5:7944295. This SNP loci is likely to influence the size of the sheep horns. To validate the identified SNP site, we employed KASP technology. Through rigorous experimental procedures, we successfully genotyped the selected SNP sites in our sample population (Fig 7B).

**Fig 7.**
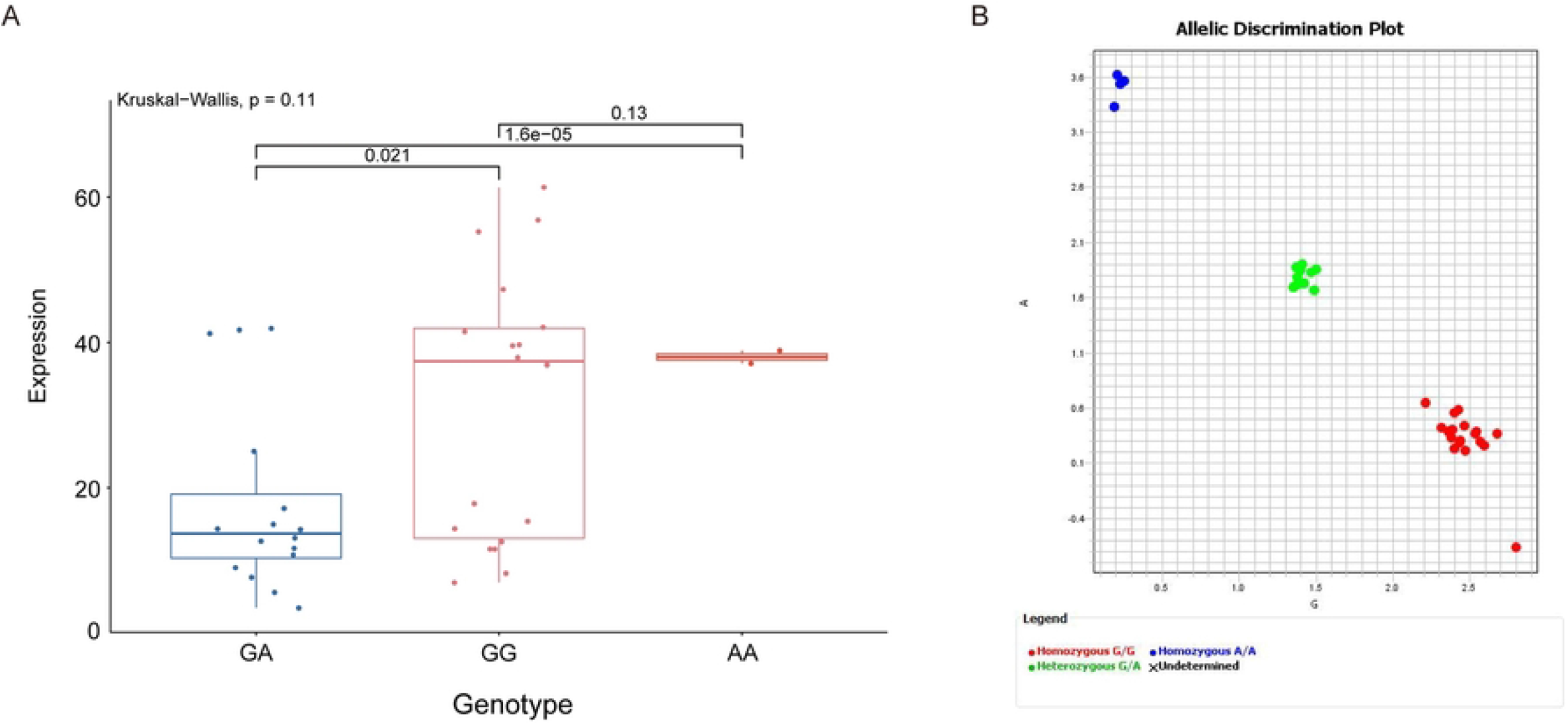
Phenotypic and Genotypic Analysis of the *CASP14* gene. (A) Box plot of individual horn lengths for different genotypes. Calculating the p-value based on the dominant genetic model and t-test. (B) The KASP assay results for the *CASP14* gene.

## Discussion

Previous studies have identified genes such as RXFP2, INSL3, KRT10, and WNT7B as being associated with horn growth [31, 32]. However, a study employing CRISPR/Cas9 to disrupt the RXFP2 gene for the first time successfully generated a cryptorchid sheep model. The findings of this study indicate that partial disruption of the RXFP2 gene did not affect the horn phenotype in sheep, suggesting that partial disruption of RXFP2 is associated with testicular descent rather than horn formation [33]. This indicates that the genes regulating the sheep horn phenotype are not singular, thus necessitating a more in-depth investigation into the genetic mechanisms underlying the sheep horn phenotype.

*CASP14* is a unique member of the evolutionarily conserved family of cysteine aspartate-specific proteases [34]. *CASP14* activation occurs between the granular and cornified layers, with both active and inactive forms detected in epidermal extracts, and only the active form in the cornified layer [35]. This activation coincides with stratum corneum formation during embryonic development [36, 37]. Our findings indicate that the *CASP14* gene is significantly upregulated in the scurred group compared to the SHE group. Additionally, the *CASP14* gene is highly expressed in the skin, exhibiting higher expression levels in both humans and sheep. The results are consistent with existing research findings [38, 39]. Our study revealed that mutations in the *CASP14* gene affect the development of skin derivatives-horn. By analyzing allele-specific expression sites (ASE), we identified five key loci within the *CASP14* gene region that may influence horn size variation. Additionally, significant differences were observed between horned and polled sheep at the screened single nucleotide polymorphism (SNP) loci. Using a dominant model and whole-genome sequencing (WGS) data, we determined that the variant locus g.7944295G>A is significantly associated with horn length in sheep. This variant likely plays a crucial role in horn growth, highlighting the impact of *CASP14* gene mutations on the morphology of skin derivatives. CK18 was expressed in normal epithelial cells of most organs but absent in normal squamous epithelium [40]. A study indicates that inhibition of *CASP14* in Be-Wo cells resulted in the upregulation of KLF4, hCG, and CK18-markers associated with normal trophoblast differentiation. These findings suggest that *CASP14* influences the differentiation pathway of trophoblasts. Our study demonstrates that the expression levels of *CASP14* are elevated in the scurred group, corroborating previous research on the influence of *CASP14* on keratinization processes.

The transcriptional regulation of the *CASP14* gene is not fully understood, with expression in keratinocytes occurring only under specific conditions like high-density culture, forced aggregation, or vitamin D3-induced differentiation [34, 38, 39, 41]. Sphingolipid metabolites, particularly ceramides, are key regulators in cellular processes and have been identified as inducers of *CASP14* [42]. The up-regulation of *CASP14* by these lipids is mediated through the inhibition of kinase pathways, highlighting its role in cell differentiation and apoptosis [43]. Treatment of human keratinocytes with Th2 cytokines IL-4 and IL-13 significantly reduces *CASP14* protein expression [44]. The *CASP14* gene is expressed in the differentiated stratum corneum and hair follicles of the epidermis, with its distribution in the epidermis and hair follicles being conserved across several mammalian species as revealed by ultrastructural analysis [45, 46]. *CASP14* is involved in the formation of the embryonic skin barrier, with its expression initiating from embryonic day 14.5 or 15.5 and processing commencing from day 17.5, coinciding with the formation of the epidermal stratum corneum [35, 47]. The predominant location of *CASP14* in the epidermis suggests it may play a specialized role there, the lowest layer of the epidermis is called the basal layer, and it consists of proliferating keratinocytes [48]. In this study, an investigation of the expression patterns of *CASP14* across various tissues revealed that the gene is highly expressed in the skin, whereas its expression in other tissues such as the heart, blood, and CNS is relatively low or nearly absent. Furthermore, the expression of the *CASP14* gene in the skin was significantly higher than in other tissues across four species. These results are similar to those of the aforementioned studies. The growth of horns is closely related to the skin, and the expression patterns of the *CASP14* protein in horned animals such as sheep, cattle, and deer within the Artiodactyla order exhibit similar trends, indicating a close association of the *CASP14* gene with horn growth.

## Conclusion

The study demonstrates a correlation between the *CASP14* gene and the size of sheep horns. Further analysis indicates that the *CASP14* gene is highly expressed in the skin tissues of both rams and ewes. We conducted an in-depth analysis of the amino acid sequence sites and protein structure of the *CASP14* gene, identifying nine amino acid positions that are specifically expressed in horned animals. Additionally, by integrating the analysis of SNPs, ASEs, and Fst of the *CASP14* gene, we identified several potential functional sites of the gene, with the SNP g.7944295 G > A potentially having a significant impact on the size of sheep horns. These sites may significantly affect the size and type of sheep horns and can serve as molecular markers for horn breeding.

## Supporting information

S1 Table. Sample infromation.

S2 Table. The mean TPM of the *CASP14* gene in various tissues of cattle, humans, pigs, and sheep.

S3 Table. SNP information of *CASP14* gene.

